# Reduced carbon emissions and chain elongation during mixotrophic fermentation of a biomass feedstock

**DOI:** 10.1101/2025.10.16.682809

**Authors:** Myra X. Afzal, Maria L. Bonatelli, Sabine Kleinsteuber, Heike Sträuber, Flávio C. F. Baleeiro

## Abstract

Anaerobic fermentation of biomass feedstocks using open cultures is a promising technology to produce platform carboxylates. Syngas, a mixture of H_2_, CO_2_, and CO, can be sourced sustainably and used to supplement biomass feedstocks as a source of acetyl-CoA, an intermediate for carboxylate chain elongation. To test this, syngas and corn silage were provided to a 10-L semi-continuous fermenter for 209 days of operation in a first-of-a-kind study at this scale. After acclimation to syngas, a two-fold reduction in average CO_2_ production rate (0.097 vs. 0.21 g L^-1^d^-1^) was observed over a period of 42 days in comparison to a control. Syngas co-feeding also increased average production rates of *n*-butyrate (C4) and *n*-caproate (C6) by 74% and 27%, respectively, although these effects were observed at relatively low C6 concentrations up to 4 g L^-1^. Relative abundances of *Megasphaera* and *Dialister* showed significant correlation (p<0.05) to consumption of H_2_ and CO as well as production of C4 and C6, suggesting involvement of these genera in mixotrophic metabolism. A feasibility analysis showed that syngas recirculation could return additional 1.54 USD m^-3^_broth_ while costing 1.26 USD m^-3^_broth_ and avoiding 1.81 kg CO_2 eq._ m^-3^_broth_ in emissions compared to heterotrophic fermentation. We propose mixotrophic fermentation as a ‘low-tech’ technology to turn fermenters into decentralized industrial carbon sinks.

**SYNOPSIS:** Fermentation of biomass and syngas could be used for sustainable and circular chemical production. This study tested the feasibility of this technology and identified relevant process parameters and microbial genera.

## INTRODUCTION

Anaerobic fermentation of abundant organic feedstocks, such as lignocellulosic biomass, offers a promising platform for the production of value-added chemicals. One such case is the conversion of agricultural residues to useful, high-value products like medium-chain carboxylates (MCCs) by anaerobic microbial communities capable of chain elongation (CE) metabolism. MCCs are monocarboxylates with linear or branched (iso) chains containing five to twelve carbon atoms in the backbone^1^. MCCs have a vast spectrum of applications in industries (e.g., cosmetics, food and feed ingredients, pharmaceuticals, and agrochemicals) and are currently produced from coconut oil, palm kernel oil, or fossil oil ^2^. Compared to other MCCs, *n*-caproate (C6) and *n*-caprylate (C8) are of particular interest because of the relatively high quantity produced by anaerobic microbial communities and the potential to be extracted by simpler and cheaper downstream processing in comparison to their short-chain counterparts^3^. Using open cultures for anaerobic fermentation reduces sterility requirements during production compared to defined cultures. It also provides a way to utilize cheap and complex feedstocks, which can be difficult for single strains to break down.

Despite these benefits, there are several challenges to overcome for fermentation of lignocellulosic biomass to become a competitive chemical production route. Organic feedstocks generally present a relative scarcity of electron donors (ED; e.g., ethanol, lactate) in comparison to electron acceptors (EA; e.g., acetate, *n*-butyrate). This scarcity of ED decreases selectivity of longer-chain, more desired chemical products ^4^ and favors competing metabolic pathways ^3, 5^. Besides, variable availability of nearby organic feedstock is one of the main bottlenecks for biorefineries to profit from economies of scale ^6^. To address these challenges, syngas, a mixture of hydrogen (H_2_), carbon dioxide (CO_2_), and carbon monoxide (CO), has been proposed as a co-substrate to microbial communities producing MCCs ^3, 7^. When syngas is co-fed with an organic feedstock, it is possible to simultaneously enrich heterotrophic and autotrophic metabolic functions in a microbial community, leading to a *de facto* mixotrophic fermentation.

H_2_ has been previously shown to be a promising ED supplement for CE when co-fed with corn silage in batch cultures ^8^, although the mechanism through which H_2_ can improve MCC production remains elusive. *Megasphaera elsdenii* has long been known as a chain elongator ^9^ that has the capability to catalyze H_2_, and the genus *Megasphaera* has been shown to be a predominant community member in systems enriched with H_2_ starting with cow manure and corn silage as inocula ^8^, making it an interesting candidate for CE in mixotrophic fermentation. CO can also act as an electron donor. Previously, syngas has been shown to be an effective ED source when co-fed with organic model feedstocks like lactate and acetate in mineral media ^10^. Moreover, CO and specific impurities commonly found in certain industrial syngas streams (e.g., ethylene, C_2_H_4_ and acetylene, C_2_H_2_) ^11^ act as methanogenesis inhibitors. For instance, the addition of CO or small doses of C_2_H_4_ has been shown to inhibit methanogenesis ^8^ and when both gases were combined, a complementary effect was observed.

Carbon fixation is another promising aspect of mixotrophic fermentation. Steering the fermentation away from CH_4_ and CO_2_ production not only re-routes electrons towards CE, but also has the potential to reduce carbon emissions of the whole process chain of a biomass feedstock. The mixotrophic and autotrophic members of a microbial community are able to utilize the CO_2_ and H_2_ produced by heterotrophic fermentation, in addition to consuming any externally supplied syngas. This syngas-consuming capability has been shown in batch cultures in several studies. ^8, 12^ The ability to steer a mixotrophic community towards syngas and synthetic growth medium consumption in a stir-tank reactor setting has also been shown ^10^. Even the external supply of CO_2_ without the other common syngas components is able to enrich some carbon-fixing functions as shown by de Leeuw, et al. ^13^ and Roghair, et al. ^14^.

The supply of CO_2_ and syngas also has implications for solventogenesis (i.e., the formation of alcohols) as part of anaerobic fermentation. Previously it was shown that some chain-elongating microbes can perform solventogenesis by reducing carboxylic acids produced in CE to their corresponding alcohols using H_2_ or CO as electron donors ^15^. Solventogenesis has been observed to be induced and tunable by parameters like pH ^16^, temperature ^17^, and CO_2_ partial pressure ^13, 14^. The potential for tunable alcohol production following CE makes solventogenesis an interesting avenue to explore in mixotrophic fermentation.

In this study, we present and evaluate the feasibility of mixotrophic fermentation with syngas and lignocellulosic biomass (corn silage, as a feedstock model) to produce MCCs in a 10-L semi-continuous fermenter. To our knowledge, this type of study has not yet been conducted at this scale. Gas composition, temperature, pH, and organic loading rate (OLR) are discussed in regard to their effects on product yield, gas consumption, carbon fixation, and MCC concentration. In addition to discussing process parameters for CE in mixotrophic fermentation and the potential of this technology as a carbon-fixing process, we also discuss the roles of key microbes such as the genera *Megasphaera* and *Dialister* in mixotrophic fermentation. The results of this study have the potential to lay groundwork towards scale-up and application of this sustainable technology.

## MATERIALS AND METHODS

### Bioreactor Operation

Two identical 15-L anaerobic stirred-tank bioreactors used previously ^18^ retrofitted with gas recirculation systems were operated for 208 days. Reactors were stirred at 150 rpm and operated at a 10-L working volume, hydraulic retention time of 10 days, and feeding every 3.5 days. Each reactor was equipped to continuously recirculate 80:20 v/v N_2_:CO_2_ (control reactor), 80:20 H_2_:CO_2_ (test reactor), or approximately 75:18:7 H_2_:CO_2_:CO (test reactor) from an 18-L flexible gas reservoir. Gas was delivered to the broth via aquarium spargers at the bottom of the reactor at a rate of approximately 1.5 L min^-1^ using a diaphragm pump (model NMP830 from KNF Neuberger GmbH, Germany) adapted with a housing for physical and electrical protection. The reactor configuration was similar to the continuous reactor described by Baleeiro, et al. ^19^. To ensure consistent recirculation of desired gases and to prevent over-pressurization due to microbial gas production, gas reservoirs were emptied using a vacuum pump and refilled every seven days with the approximate compositions described above. The partial pressures of non-inert gases across the experiment are shown in Figure S1. The gas balance in the first 70 days was not used to show gas accumulation data because the He concentrations were sometimes too low and the data were not reliable. Presence of N_2_ was used as an indirect measurement of O_2_ contamination from air. This method to estimate O_2_ contamination has been described in a similar system before ^20^. For detailed information on reactor operation, please refer to the Supporting Information.

Reactors were operated at 32°C and pH 5.5, with the exception of days 146-181, when pH and temperature alternated between 32°C/pH 5.5 and 28°C/pH 4.8 every 3.5 or 7 days. A detailed description of reactor operation phases and process conditions can be found in Table S1. OLR was maintained at either 3.6 or 7.2 g volatile solids (VS) L^-1^ d^-1^ with a constant hydraulic retention time of 10 d. These two OLR values were named “low” and “high”, respectively. Corn silage from Neichen, Germany, was used as substrate, and sludge from a mesophilic biogas reactor fed with corn silage and cow manure operated by the *Deutsches Biomasseforschungszentrum - DBFZ* (Leipzig, Germany) was used as inoculum. The inoculum and the corn silage were sampled in triplicate to determine total solids (TS), VS, ammonia nitrogen content (NH_4_-N), and pH, and the means of these values are reported in Table S2. The microbial composition of the inocula is presented in Figure S2.

### Sample Collection and Chemical Analysis

Gas composition (helium - He, hydrogen - H_2_, carbon dioxide - CO_2_, ethylene - C_2_H_4_, methane - CH_4_, oxygen - O_2_, nitrogen - N_2_, and carbon monoxide - CO) was monitored five times per week using two gas chromatography (GC) systems (MicroGC Fusion, Inficon, Germany; PerkinElmer, USA). Analysis by PerkinElmer GC was done as described by Logroño, et al. ^21^. Analysis by MicroGC was performed as described by Mohr, et al. ^22^, where the sampling volume was 2 mL. Broth was sampled from the reactor harvest prior to feeding, which occurred twice a week. Broth was analyzed for pH, TS, VS, ammonia nitrogen, and organic acids and alcohols content. TS and VS analyses were done as described by Strach ^23^. Ammonia nitrogen content was measured as described by Strach ^24^.

Chemical content of the broth was measured by high-performance liquid chromatography (HPLC) and by a GC system coupled to a flame ionization detector with samples preceding a derivatization by esterification (ester-GC) adapting methods from Apelt ^25^ and Apelt ^26^, respectively. Modifications to these methods are described in the Supporting Information. Measured compounds include formate (C1), acetate (C2), lactate (LAC), ethanol (EtOH), propionate (C3), *n*-propanol (PropOH), *n-*butyrate (C4), *i*-butyrate (iC4), *n*-butanol (ButOH), *n*-valerate (C5), *i-*valerate (iC5), *n*-pentanol (PentOH), *n-*caproate (C6), *i-*caproate (iC6), *n*-hexanol (HexOH), *n-*heptanoate (C7), and *n-*caprylate (C8).

Rates in this study are average rates of production or consumption specific to the broth volume (10 L). Average rates calculated in mmol of electron equivalents (e^−^ mmol) were used to compare chemicals in the gas and liquid phases. The conversion factors for realizing molar, electron, carbon and material balances are presented in Table S3. The original experimental data and calculated rates are included in the Supporting Information.

### Microbial Community Analysis

Twice a week, 5.4 mL of broth was collected for microbial community analysis. Centrifuged cell pellets free of supernatant were resuspended in 1 mL of 10 mM Tris-HCl buffer (pH 8.5) and centrifuged for 10 minutes at 20,000 × g. Supernatant was removed and the pellets were stored at -20°C for microbial community analysis.

DNA was extracted using the NucleoSpin Soil Kit (Macherey-Nagel, Germany) following the manual’s instructions and DNA quantification was conducted with Qubit™ dsDNA BR Assay Kit (Invitrogen, Germany). Amplicons covering the V3-V4 region of the 16S rRNA gene were generated using the primer set 341f (5’-CCT ACG GGN GGC WGC AG-3’) and 785r (5’-GAC TAC HVG GGT ATC TAA TCC-3’) ^27^. The library was prepared with MiSeq Reagent Kit v3 with 2 × 300 cycles and sequenced on the MiSeq platform (Illumina, San Diego, CA, USA).

Amplicon sequence analysis was done according to Baleeiro, et al. ^28^. Briefly, Cutadapt ^29^ was used to remove primer sequences, and DADA2 to infer amplicon sequence variants (ASVs) ^30^. Phyloseq ^31^ was used for diversity analysis in R R Core ^32^. All samples were rarified to an equal sequencing depth of 6730 counts. Spearman’s correlation analysis was done between the relative abundance of ASVs, after a centered log-ratio transformation to account for compositionality ^33^, and the concentrations of the different organic chemicals measured. Raw sequence data was deposited at the European Nucleotide Archive (ENA) under the study accession number PRJEB88737 (http://www.ebi.ac.uk/ena/data/view/PRJEB88737).

### Techno-economic Analysis

Detailed assumptions adopted in the techno-economic analysis are presented in the Supporting Information.

## RESULTS AND DISCUSSION

Reactor operation for this study was performed semi-continuously for 208 days. The experiment can be divided into five phases that compared H_2_:CO_2_ vs. N_2_:CO_2_ and syngas vs. N_2_:CO_2_, each at high and low OLR, as well as varying pH and temperature. The reactors were restarted once at day 104. Representations in chemical concentrations, absolute quantities, as well as a detailed description of reactor conditions are provided in the Supporting Information.

During reactor startup (days 0-70), reactors operated with N_2_:CO_2_ or H_2_:CO_2_. Chemical concentrations were similar between the control and test reactors (Figure S3). The results for the startup are presented and discussed in depth in the Supporting Information.

### H_2_ and C_2_H_4_ Shift Community Composition

During the first 70 days, both reactors showed similar chemical compositions (Figure S3). Measurements of gas composition during this period were unreliable and are not reported. Under high OLR (7.2 g VS L^-1^d^-1^, days 0 to 28), H_2_ did not appear to impact microbial composition. Overall, differences in the chemical and community compositions between the reactors with and without added H_2_ in the first 70 days had very similar trends (Figure S3). The similarity between the two reactors in the first 28 days could be attributed to the small difference in H_2_ partial pressure between control (7.3 kPa) and test (17.6 kPa) reactors. The relatively low H_2_ partial pressure in the test reactor for this period was due to faulty gas bag replenishment, which was later corrected. With the increase in H_2_ supply after day 26, methanogenesis took off and rapidly consumed most of the H_2_ supplied to the test reactor (Figure S1 A).

C_2_H_4_ was added to the test reactor on day 49 and had an immediate effect of inhibiting methanogenesis and consequently guaranteeing an increased H_2_ partial pressure (Figure S1 A). Despite the increased H_2_ partial pressures after C_2_H_4_ addition (average of 33.0 kPa for the whole period in comparison to 10.7 kPa in the control), only minor changes in the chemical concentrations of the test reactor were observed, such as a slight increase of concentration of C2 and C4 and a decrease of C8 concentration (Figure S3). Under increased H_2_ partial pressure, genera such as *Prevotella, Syntrophococcus*, and *Dialister* had higher relative abundance compared to the control, while the relative abundance of *Lactobacillus* decreased under H_2_ conditions (Figure S3).

Figure 1 presents the electron balance for the period between 70 and 91 d. Here, successful suppression of methanogens in the test reactor allowed for bacterial H_2_ consumers to emerge as net H_2_ consumption in the test reactor could be observed, despite having lower methanogenic activity and the control showing a net production of H_2_ and a higher CH_4_ production (Figure 1). Still, the amount of H_2_ consumed at this stage was not enough to impact chemical accumulation. With the exception of *Syntrophococcus* and *Solobacterium*, the community composition (e.g. the order of genera abundance) of both reactors showed very similar trends as seen in Figure S3 for the 70 – 91 d period.

**Figure 1.**
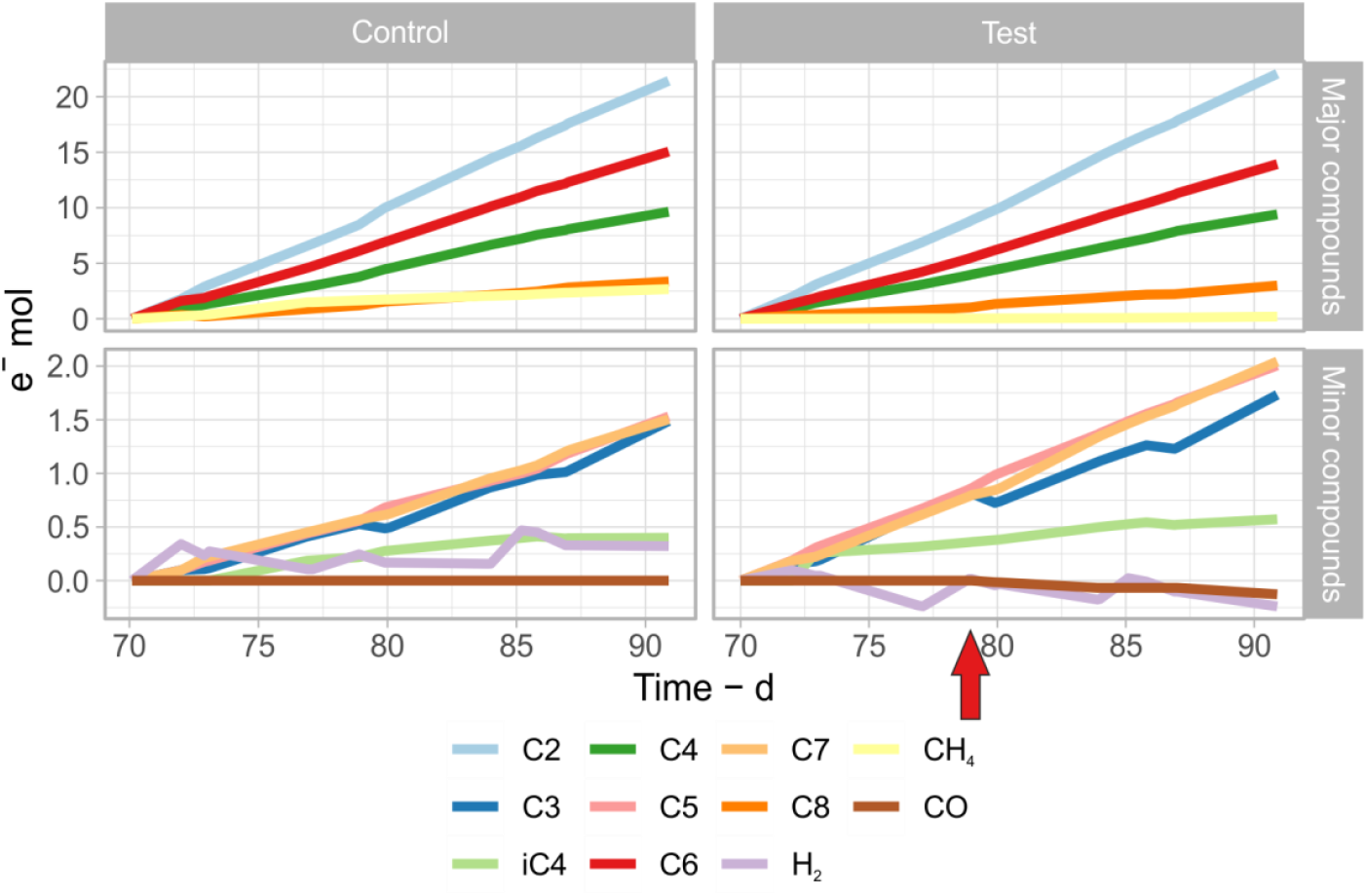
Accumulation of mol-electron equivalents over time for the most relevant gases and carboxylates between days 70 and 92. C_2_H_4_ was used as a methanogenesis inhibitor in the test reactor at this time; the red arrow on day 79 indicates the first CO addition to the test reactor. Gas balances prior to day 70 are not shown because tracer gas concentrations were not reliable enough to calculate accurate balances.

### CO Addition and OLR Likely Influence H_2_ Consumption

CO started to be injected on day 79 and initially maintained an average partial pressure of 2.0 kPa in the test reactor (Figure S1 A). An incipient consumption of CO is visible starting around day 86, about 7 days after its first addition, which indicates a short acclimation period.

At this point of the experiment, several sparging stones responsible for gas delivery into the reactor came out during the reactor harvest, leading to poor gas delivery. To fix this, the reactors were emptied on day 98 and more robust sparging stones were adhered to the bottom of both reactors (Figure S4) before their restart on day 104. Figure 2 shows the concentration of main chemicals and the community composition after this restart, which occurred under low OLR (3.6 g VS L^-1^d^-1^) and a complete syngas mixture (i.e., H_2_, CO_2_, CO and C_2_H_4_) in the test reactor.

**Figure 2.**
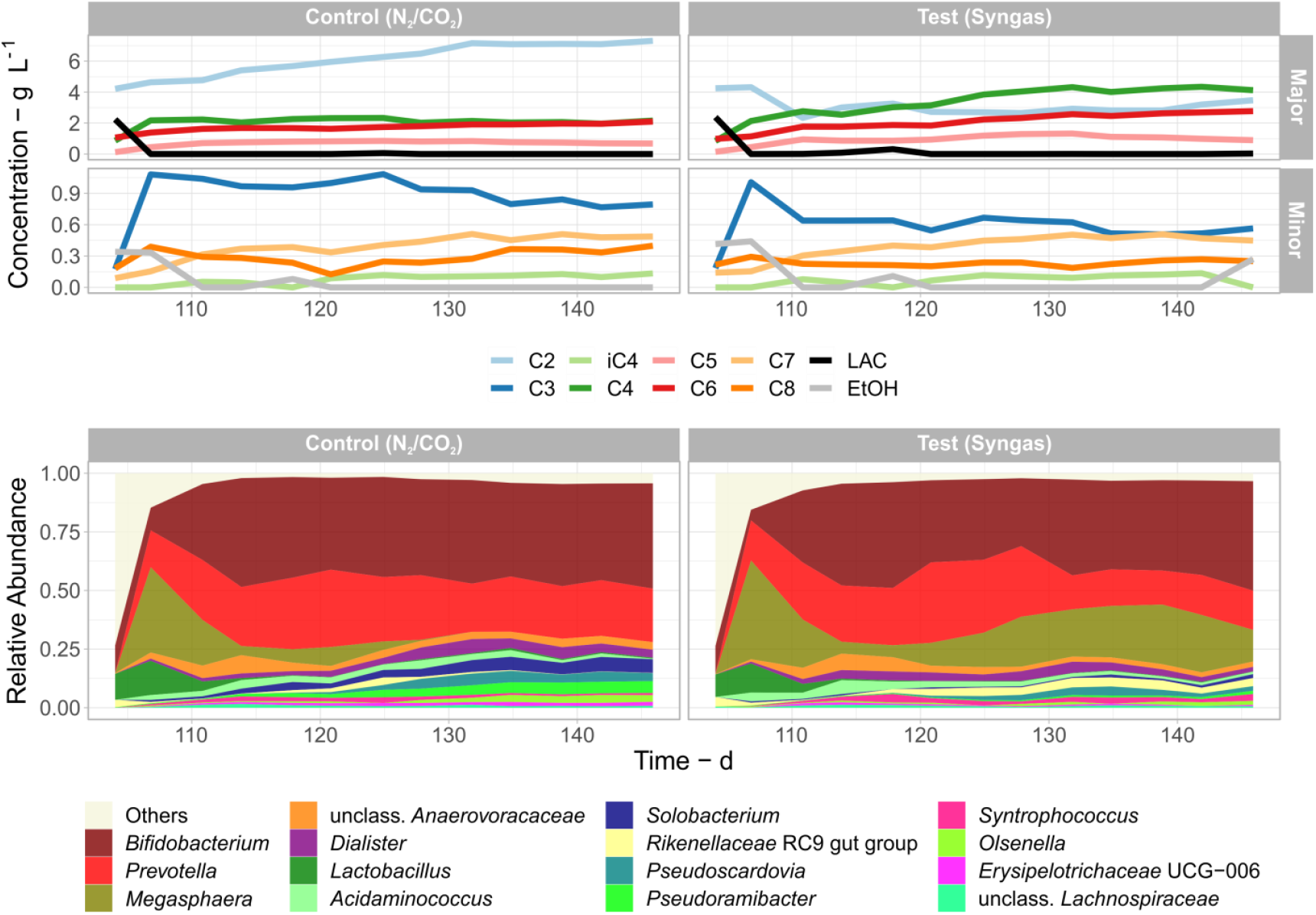
Chemical concentrations (top) and the top 15 most abundant genera present (bottom) over time when both reactors were operated at OLR=3.6 g VS L^-1^ d^-1^ and the test reactor was operated with syngas.

Under low OLR, C4 and C6 started to be produced almost immediately in both the control and test reactors, although the test reactor showed generally higher concentrations of C4 and C6 and lower C2 and C3 concentrations than control (Figure 2). *Megasphaera* was the key differentiating genus in the test reactor with syngas, which was not observed during the initial reactor stages. After an initial increase in both reactors, *Megasphaera* was outcompeted in the control reactor. In the test reactor, *Megasphaera* remained one of the three dominant genera, matching with increased C4 and C6 production (Figure 2).

The effect of syngas was also tested under high OLR (7.2 g VS L^-1^d^-1^) between days 180 and 208. The resulting community structure and chemical production are shown in Figure S5. Under high OLR, both reactors showed similar chemical and community profiles, with slightly higher C2 and C3 concentrations, lower relative abundances of *Olsenella* and *Caproiciproducens*, and a transient LAC accumulation in the test reactor. *Megasphaera* was not among the 15 most abundant genera from the beginning on day 181 until the end of this period on day 208, although it is not clear if its loss of relevance in the community was due to the high OLR, as seen in previous studies ^34^, or due to the pH cycling of the preceding experimental phase (Figure S6). Despite the washout of *Caproiciproducens* in the test reactor, C6 concentration in both reactors was similar, suggesting that *Pseudoramibacter* was the main producer of longer-chain carboxylates under high OLR (Figure S6).

The average molar production rates for the main chemicals are presented for both low and high OLR conditions in Figure 3. Under these conditions, the average C4 and C6 production rates in the test reactor were 4.72 and 2.16 mmol L^-1^d^-1^, respectively, representing a 74% and 34% increase compared to the control over a 42-day period (Figure 3B). The increased MCC production in the test reactor coincided with H_2_ and CO consumption (9.35 mmol L^-1^d^-1^ and 0.88 mmol L^-1^d^-1^, respectively), suggesting that H_2_ and CO consumption was important for C4 or C6 production. Net H_2_ consumption did not occur in the control reactor.

**Figure 3.**
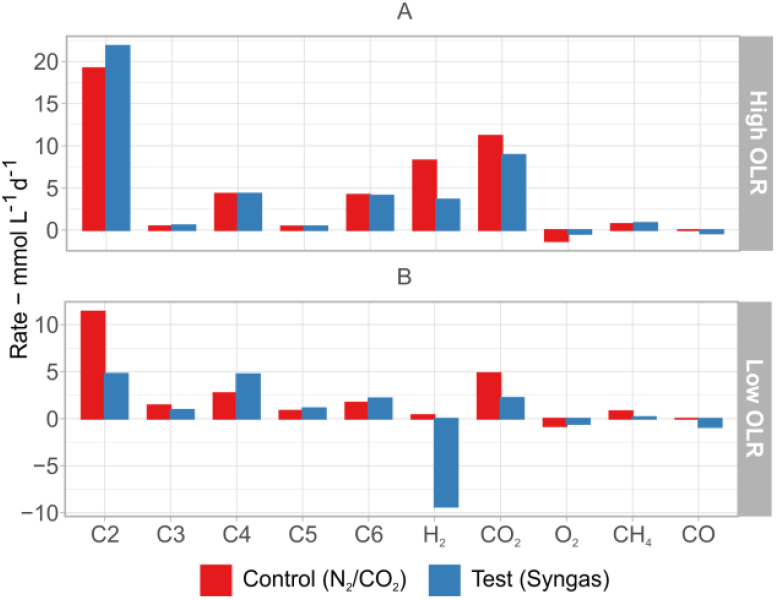
Average production and consumption rates for select chemicals and gases between days 180-208, OLR=7.2 g VS L^-1^d^-1^ (A) and days 104-146, OLR= 3.6 g VS L^-1^d^-1^ (B). Both reactors were operated with syngas during these periods. Negative rates denote consumption.

Given the lack of H_2_ consumption prior to the addition of CO in the test reactor, and little to no H_2_ consumption in the control reactor, CO presence was the defining factor to induce H_2_ consumption and subsequent C4 production and chain elongation (i.e. C6 production).

### pH and Temperature Modulation Lead to Decreased *Megasphaera* Abundance

To induce solventogenesis with CO and H_2_, pH and temperature were adjusted every 2.8 or 6.8 days between 32°C/pH 5.5 and 28°C/pH 4.8. PropOH and ButOH were used as solventogenesis indicators, but their concentrations remained minimal (<0.2 g L^−1^). Chemical concentrations were very similar in both the control and test reactors, pointing out that the reducing power of CO and H_2_ was not used to reduce carboxylates into alcohols to a relevant extent. Figure S6 presents the time profiles for chemical concentration, community composition, and pH variation for this experiment. An in-depth discussion about this experiment can be found in the Supporting Information.

### Community Members Involved in Syngas Consumption and CE

According to Spearman correlations (Figure 4), relative abundances of the ASVs *Megasphaera* (008), *Prevotella* (007), *Syntrophococcus* (017), and unclassified *Lachnospiraceae* (020) were significantly (p<0.05) correlated to H_2_ consumption, whereas *Dialister* (015), *Prevotella* (012), and *Syntrophococcus* (014) correlated significantly with CO consumption. In terms of CO_2_ consumption (or lower CO_2_ emissions), *Bifidobacterium* (002), *Dialister* (015), and *Streptococcus* (017) presented the only significant correlations (Figure 4). Among the ASVs correlated with consuming gases, *Megasphaera* (008), *Prevotella* (012), and *Dialister* (015), were correlated to C4, C5, and C7 concentrations in varying degrees (Figure 4). Additionally, *Pseudoramibacter* (005) was recognized as a heterotrophic chain elongator, effectively producing C6 and C8 fatty acids without consuming H_2_, CO, or CO_2_.

**Figure 4.**
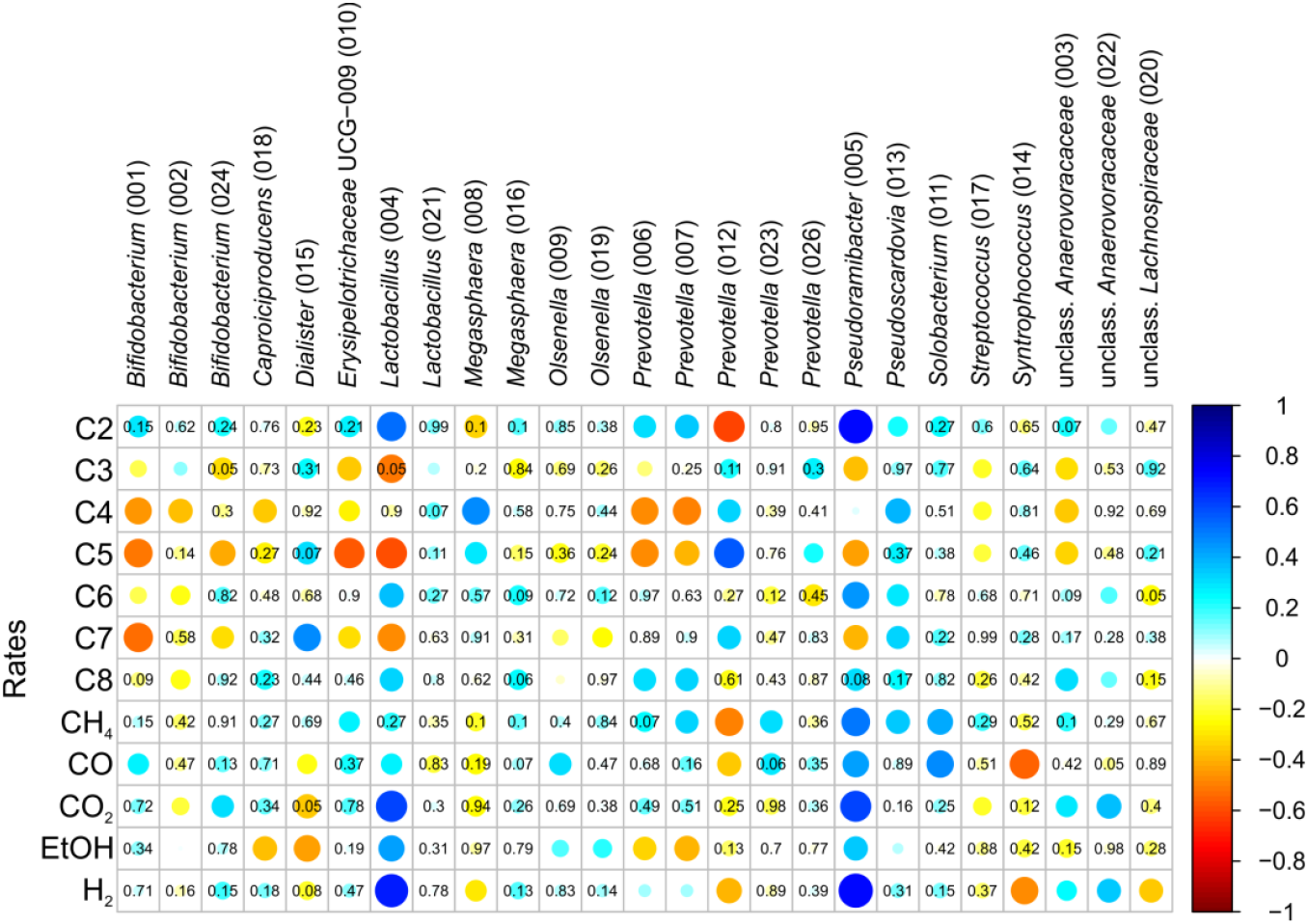
Spearman correlation between abundance of ASVs and production/consumption of selected chemicals across the entire experiment. Blue shades indicate a positive correlation, showing that the ASV was abundant when the chemical production rate was high or when the consumption rate was low. Red shades indicate a negative correlation. Larger circles with darker shades indicate a stronger correlation between ASVs and chemicals. p-values are only shown for the less significant correlations (p≥0.05). The relative abundances were corrected for compositionality with a centered-log ratio transformation before calculating correlation coefficients.

Both *Megasphaera* and *Dialister* belong to the *Veillonellaceae*. While we previously discussed the overlooked potential of *Megasphaera* to consume H_2_ and grow mixotrophically^9, 28^, no carboxydotrophic members of this family are known. Still, many putative carbon monoxide dehydrogenase (CODH) genes have been found in genomes affiliated to the *Veillonellaceae*^35^. The UniProt database^36^ returns 123 entries for CODHs and homologue proteins (e.g., hydroxylamine reductase) predicted or inferred from *Veillonellaceae* genomes (as of 04/28/2025 with the keyword “CODH”). These CODH genes are often accompanied by gene sets of the Wood-Ljungdahl pathway^35^. Even though many of these gene sets lack the gene for formate dehydrogenase^35^ used to convert CO_2_ into formate, formate is a common intercellular metabolite, and we have previously shown its importance as an electron and carbon carrier between bacteria in mixotrophic fermentation of biomass and syngas ^8^. Therefore, we suspect that *Megasphaera* and *Dialister* compose the main mixotrophic group in our system. It is also possible that both community members have different gas-consuming pathways despite their taxonomic relation. In open culture reactors without syngas addition, *Dialister* abundance has been shown to decrease when CO_2_ supply was removed^37^, which could also point to a CO-independent carbon fixation mechanism in it.

There were two other candidate mixotrophic groups in our system: *Prevotella* (012) and the two *Lachnospiraceae* ASVs: unclassified genus (020) and *Syntrophococcus* (014) (Figure 4). Like the *Veillonellaceae*, many members of the *Lachnospiraceae* have recently been found to harbor genes for the Wood-Ljungdahl pathway^35^. However, the *Lachnospiraceae* ASVs had very low relative abundances in this study and while they correlated significantly with H_2_ and CO consumption, we could not associate them with any carboxylate product (Figure 4). For the *Prevotella* ASV that presented some correlation with syngas components, an indirect correlation might be the best explanation. The lactic acid bacteria and hydrolytic groups, including *Bifidobacterium, Lactobacillus*, and *Prevotella*, are likely crucial for supplying intermediates to other acidogenic bacteria and should be particularly prone to cross-correlation with diverse metabolic functions. Besides, we previously observed and discussed these possible cross-correlations between H_2_ and CO consumption and lactic acid bacteria ^8^. For *Prevotella*, a search for CODH in UniProt only returns genes for homolog enzymes (*hcp* or hydroxylamine reductases).

*Megasphaera* was favored when enriched with H_2_, CO, and educts from lignocellulosic biomass as EDs (Figure 2). The isolation paper of *Megasphaera elsdenii* has shown its capabilities of consuming H_2_ and producing a wide range of carboxylates from C3 to C6 ^38^. However, to our knowledge, the potential of this genus to adopt a stable mixotrophic role in an open culture has not been shown previously. The dominance of a known chain elongator genus such as *Megasphaera* in a complex system, such as the one in this study, hints at ubiquitous potential of enriching mixotrophic open cultures that are industrially relevant. If mixotrophic communities proof as easy to enrich such as here, there is an untapped potential of using syngas in biomass fermentation for forming microbial communities that valorize C1 compounds and H_2_ into economically attractive products.

### Factors for Reduced Carbon Emissions

One interesting outcome of this experiment was reduced CO_2_ emissions from the test reactor under mixotrophic conditions. By summing the molar production rates of CH_4_, CO_2_, and CO in Figure 3, a value for molar carbon emissions can be obtained. Average C emission rates in C mmol L^-1^d^-1^ calculated for all experimental periods are available in the spreadsheet file provided in the Supporting Information.

Under low OLR (3.6 g VS L^-1^d^-1^), the test reactor emitted on average 73% less carbon (1.50 C mmol L^-1^d^-1^ compared to 5.61 C mmol L^-1^d^-1^ in the control). In parallel, it produced 74% more C4, 23% more C6, and 79% less CH_4_ than the control reactor (Figure 3B). Under high OLR (7.2 g VS L^-1^d^-1^), the reduction of carbon emissions was 21% (9.37 C mmol L^-1^d^-1^ compared to 11.9 C mmol L^-1^d^-1^ in the control), presenting a tradeoff between biomass utilization and carbon emissions.

Regarding MCC production, at low OLR, relatively low MCC concentrations (2.2 g C6 L^-1^) were sustained in comparison to 4.3 g C6 L^-1^ achieved at higher OLR. In recent literature, more than 10 g C6 L^-1^ has been reported under optimal chain elongation conditions ^39^. Despite the benefit of higher MCC concentration associated with high OLR, the reduced carbon emissions at low OLR should not be overlooked. Steady-state low MCC concentrations are not a general obstacle, as several in-line extraction technologies exist that are currently being scaled up ^40^ and would overcome this potential drawback.

It is important to note that the maximum syngas consumption rate in this study was likely below its maximum capacity due to relatively low partial pressures of H_2_ and CO (Figure S1A). The gas reservoirs used here were relatively small in comparison to the reactor volume (1.8:1 gas-to-reactor volume ratio) causing H_2_ and CO partial pressures to drop quickly in case of consumption or dilution with biogas and inert gases. Higher gas-to-reactor volume ratios and more frequent refilling procedures will facilitate fermentation with higher syngas consumption activity. Therefore, we assume that a fermentation with a net carbon fixation balance and a stronger effect of syngas on the carboxylate production is technically feasible.

The ability of a mixotrophic fermentation process to simultaneously utilize syngas and reduce CO_2_ emissions compared to fermentation without syngas has significant implications for the potential of this technology to reduce the environmental impact. One possibility is the creation of decentralized industrial carbon sinks by low-tech conversion of anaerobic fermenters into mixotrophic gas fermenters.

### Turning Fermentation into an Industrial Carbon Sink

In comparison to a conventional anaerobic bioreactor that only ferments biomass, a mixotrophic operation adds three major layers of operational complexity: 1) H_2_ and CO have to be supplied or generated in situ; 2) continuous gas recirculation means that a blower or compressor needs to be employed; and 3) inert gases (e.g., N_2_ and CH_4_) need to be removed from the recirculating gas to avoid indefinite accumulation. Each of these points is associated with capital costs, operational costs and environmental impacts. Here, we try to gauge the size of operational costs and emissions associated with points 1) and 2).

To calculate the costs associated with point 2), a range of the main factors for compressor electricity consumption (i.e., pressure increase and gas flow) has to be defined. In this study, a constant gas recirculation rate of 9.0 ‘reactor volumes’ per hour (9.0 vvh, in h^-1^, equivalent to 0.15 vvm, in min^-1^) and an average pressure increase of 70 mbar were used. These values help set the upper and lower limits for gas flow rate and pressure, respectively. For reference, the gas flow is 85,000-fold higher than the summed H_2_ and CO microbial consumption rates observed at low OLR (252 mL d^-1^ L^-1^ or 1.05×10^-4^ vvh). Figure 5 presents the expected operating costs and associated carbon emissions due to continuous gas recirculation rates from 1.05×10^-4^ to 9.0 vvh and compressor pressure increase from 70 to 2000 mbar. The expected operating costs of acquiring H_2_ and CO at spot market prices are shown in Figure 5A, while the range of avoided carbon emissions in kg CO_2 eq._ m^-3^ broth are shown in Figure 5B. The full set of assumptions adopted for these estimates is presented in the Supporting Information.

**Figure 5.**
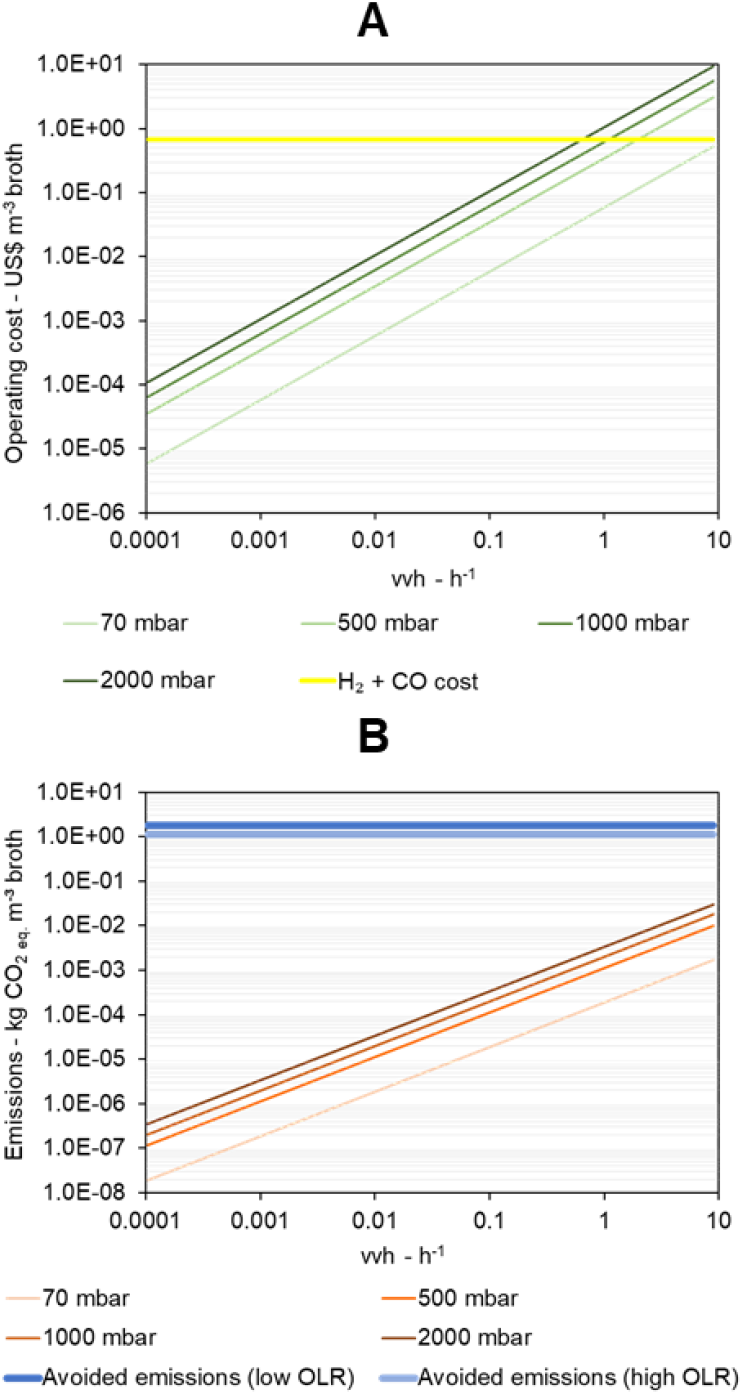
Operating costs of recirculating gas in USD m^-3^_broth_ (A) and carbon emissions in kg CO_2 eq._ m^-3^_broth_ (B) due to compressor electricity consumption depending on gas flow rate intensities in reactor volumes per hour (vvh) and pressurization needed.

According to Figure 5A, gas flow rates between 0.5 and 9 vvh are well within the operational range of industrial scale mixotrophic fermentation and would incur in operating costs that are in the same order of magnitude as the costs to supply blue or green H_2_ and CO, which are expected to be around 0.66 USD m^-3^_broth_, depending on the compressor pressurization. On the other hand, decreasing gas flow rates to below 0.1 vvh would result in gas recirculation costs that are orders of magnitude lower than the costs for supplying H_2_ and CO. However, we assume that using such low gas flow rates without decreasing microbial gas consumption rates would be technically challenging.

The impact of gas recirculation on the carbon emissions should be framed differently than its impact on costs. Recirculating gas would be associated with carbon emissions no greater than 0.04 kg CO_2 eq._ m^-3^_broth_, which is well below the least of avoided C emissions that mixotrophy can provide (between 1.11 and 1.81 kg CO_2 eq._ m^-3^_broth_ according to our experiment) (Figure 5B). This is true even considering that our assumptions adopt a very conservative electricity emission intensity of 0.481 kg CO_2 eq._ kWh^-1^, which is about half the emissions of coal-generated electricity. Since the avoided emissions due to microbial activity greatly outpaces emissions due to electricity consumption, ensuring efficient gas recirculation is a critical factor in terms of operating costs rather than in terms of carbon emissions.

Figure 6 presents possible operating costs for mixotrophic operation by combining gas circulation rates, compressor pressurization, and H_2_ and CO market prices to generate three scenarios with increasing techno-economic difficulty at commercial scale. Scenarios 1, 2, and 3 represent operating costs based on varying gas acquisition and electricity costs and are further described in the Supporting Information. These scenarios correspond to costs of 0.498, 1.26, and 2.11 USD m^-3^_broth_, respectively, and can be compared with possible broth values ranging from 7.0 to 14.4 USD m^-3^_broth_, depending on the operating conditions (Figure 6). In all scenarios, the value of the broth surpasses the operating costs. Operating the fermenter mixotrophically valorizes 1.40 USD m^-3^_broth_ by increasing C4 and MCC content and an extra 0.14 USD m^-3^_broth_, if carbon credits can be accounted for. At low OLR, mixotrophic fermentation could support a total broth value of 8.54 USD m^-3^_broth_. Assumptions for Figure 6 are described in the Supporting Information. Table S4 presents the extraction efficiencies for different carboxylates, based on Braune, et al. ^41^, and the assumed carboxylate selling prices used to estimate broth value.

**Figure 6.**
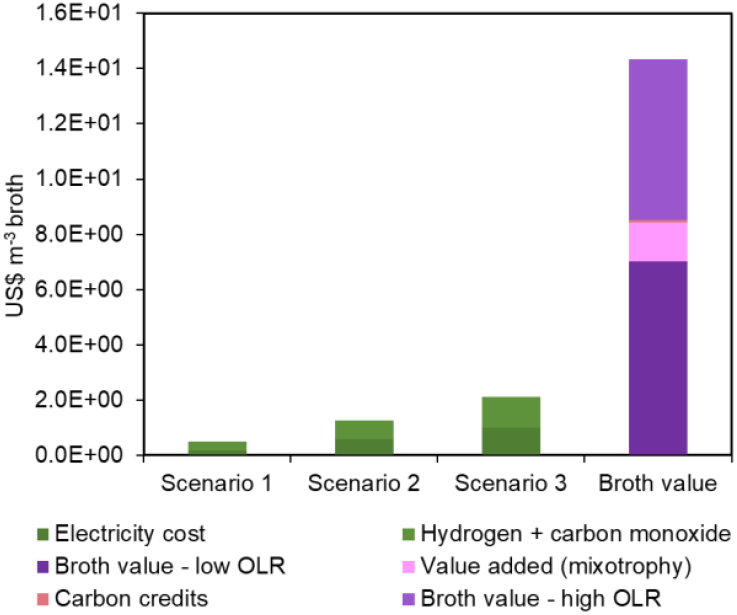
Comparison between the operating costs for supplying H_2_ and CO and continuously recirculating gas compared to the value of the carboxylates in the fermentation broth on a m^3^ of broth basis.

It is important to note that the point of this feasibility analysis is to compare mixotrophic operation with a conventional carboxylate fermentation without syngas fermentation. In other words, all operating costs that incur in conventional carboxylate fermentation (e.g., feedstock procurement, chemicals, and downstream processing costs) were not included here. Despite this, this study shows that mixotrophic fermentation has the potential of not only being environmentally favorable – by turning carboxylate fermenters into carbon sinks – but also economically viable. The techno-economic scenarios 1 and 2, which would support the break-even of gas recirculation, are well within realistic boundaries since they assume blue or green H_2_ prices not higher than 3 USD kg^-1^, gas flow rates of up to 1.0 vvh, and compressor pressurizations of up to 1000 mbar (see Supporting Information for more details).

We recommend further research of mixotrophic fermentation that deals with its current technical shortcomings and unknowns. In particular, solutions that 1) decrease the gas recirculation rates without decreasing microbial gas consumption, 2) increase the concentration of extractable carboxylates in mixotrophic communities, 3) sustain microbial gas consumption under higher OLR, and 4) investigate the equipment and installations needed for commercial scale gas recirculation.

## Supporting information

Raw experimental data and calculated average production rates

## ASSOCIATED CONTENT

### Supporting Information

Additional figures, tables, and text describing reactor operation conditions, reactor configuration, chemical concentrations, analyzed microbial data, and calculations for economics and mass balances (PDF).

Raw experimental data and calculated average production rates (XLSX).

## AUTHOR INFORMATION

### Author Contributions

MXA, SK, HS, and FCFB conceptualized the study. MXA, MLB, SK, HS, and FCFB reviewed the manuscript. MXA, MLB, and FCFB developed the methodology, analyzed the data, and developed the visualizations. MXA performed the experiments and prepared the original draft. SK, HS, and FCFB supervised the project. All authors have given approval to the final version of the manuscript.

### Funding Sources

Financial support was received from the Deutscher Akademischer Austauschdienst (DAAD) through the One-Year Grants for Doctoral Candidates Research Grant, 2020/21 (Reference 57507870), from the European Union as part of the ‘Saxony-Anhalt Science Education, Research and Development (JTF)’ program of the state of Saxony-Anhalt (Project NWG@HoMe, funding code ZS/2023/12/182203), and from the Helmholtz Association (Research Program Earth and Environment).

### Notes

The authors declare no competing financial interests.

## ACKNOWLEDGMENT

We thank Ute Lohse for technical assistance in library preparation for MiSeq amplicon sequencing and Jana Raab for assistance with reactor operation and sample collection. The DBFZ Department Biochemical Conversion is acknowledged for providing the reactors and support for the ester-GC, TS, VS, and NH_4_-N analysis. We thank the Deutscher Akademischer Austaschdienst (DAAD) for funding the one-year research exchange necessary to carry out this project. The graphical abstract was created in BioRender. Bonatelli, M. (2025) https://BioRender.com/fowz2gt.

## ABBREVIATIONS

MCC: medium-chain carboxylate;
CE: chain elongation
EA: electron-acceptor
ED: electron-donor
HPLC: high-performance liquid chromatography
VS: volatile solids
TS: total solids
GC: gas chromatography
OLR: organic loading rate
ASV: amplicon sequence variant
C1: formate
C2: acetate
EtOH: ethanol
LAC: lactate
C3: propionate
PropOH: 1-propanol
C4: *n-*butyrate
iC4: *i-*butyrate
ButOH: 1-butanol
C5: *n-*valerate
iC5: *i-*valerate
PentOH: 1-pentanol
C6: *n-*caproate
iC6: *i-*caproate
HexOH: 1-hexanol
C7: *n-*heptanoate
C8: *n-* caprylate
CODH: carbon monoxide dehydrogenase.

---

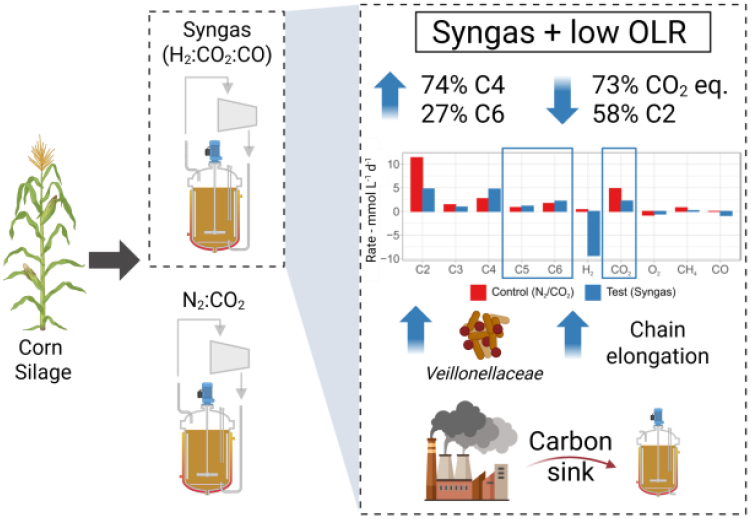

For Table of Contents Only. Created in BioRender. Bonatelli, M. (2025) https://BioRender.com/fowz2gt

## Notes

http://www.ebi.ac.uk/ena/data/view/PRJEB88737

### Competing Interest Statement

The authors have declared no competing interest.

