## Supporting Information for "Reduced carbon emissions and chain elongation during mixotrophic fermentation of a biomass feedstock"

**Table S1.** Reactor operation timeline.

| *Day* | *OLR (g VS $L^{-1}d^{-1}$)* | *pH* | *Temperature (℃)* | *Notes* |
|---|---|---|---|---|
| *0 – 28* | 7.2 | 5.5 | 32 | Inoculation of both reactors on day 0.<br>The test reactor operated with $H_2/CO_2$.<br>Not enough He added to the systems for estimating mass balances. |
| *29 – 91* | 3.6 | 5.5 | 32 | Inoculation on day 0.<br>Day 49: $C_2H_4$ added to test reactor.<br>Day70: sufficient He added to both reactors for mass balances.<br>Day 79: CO added to test reactor. |
| *91 – 98* | 3.6 | See notes | See notes | First attempt at a pH cycling experiment. pH control tuned off (pH ~4.5) and 28°C.<br>Results not shown due to problems with the gas sparging system. |
| *104 – 146* | 3.6 | 5.5 | 32 | Re-inoculation of both reactors on day 104.<br>Test reactor operated with syngas ($H_2$ + $CO_2$ + CO). |
| *146 – 166* | 3.6 | See notes | See notes | Alternate between 32°C/pH 5.5 and 28°C/pH 4.8 every 3.5 days; test reactor operated with syngas. |
| *166 – 180* | 3.6 | See notes | See notes | Alternate between 32°C/pH 5.5 and 28°C/pH 4.8 every 7 days; test reactor operated with syngas. |
| *180 – 208* | 7.2 | 5.5 | 32 | Test reactor operated with syngas. |

**Table S2.** Corn silage and inoculum characteristics, including pH, ammonia nitrogen content ($N\text{-}NH_4$), percent total solids (TS %), and percent volatile solids (VS %).

| *Sample* | *pH* | *N-NH$_4$* (g L$^{-1}$) | *TS* (%) | *VS* (%TS) |
|---|---|---|---|---|
| Corn silage 1 | 4.02 | 0.35 | 38.20 | 96.87 |
| Corn silage 2 | 3.92 | 0.44 | 32.48 | 97.35 |
| Inoculum day 0 | 7.68 | 1.75 | 2.45 | 74.19 |
| Inoculum day 104 | 7.69 | 1.56 | 2.99 | 76.34 |

**Table S3.** Conversion factors for electron and carbon balances.

| *Compound (formula)* | *Molar mass* (g mol$^{-1}$) | *mol C/mol* | *mol e$^-$/mol* |
|---|---|---|---|
| C1/Formic acid/formate ($CH_2O_2$) | 46.00 | 1.0 | 2.0 |
| C2/Acetic acid/acetate ($C_2H_4O_2$) | 60.02 | 2.0 | 8.0 |
| EtOH/Ethanol ($C_2H_6O$) | 46.02 | 2.0 | 12.0 |
| C3/Propionic acid/propionate ($C_3H_6O_2$) | 74.03 | 3.0 | 14.0 |
| PropOH/Propanol ($C_3H_8O$) | 60.09 | 3.0 | 18.0 |
| LAC/Lactic acid/lactate ($C_3H_6O_3$) | 90.03 | 3.0 | 12.0 |
| C4/Butyric acid/butyrate ($C_4H_8O_2$) | 88.04 | 4.0 | 20.0 |
| iC4/*i*-Butyric acid/*i*-butyrate ($C_4H_8O_2$) | 88.04 | 4.0 | 20.0 |
| ButOH/Butanol ($C_4H_{10}O$) | 74.04 | 4.0 | 24.0 |
| C5/Valeric acid/valerate ($C_5H_{10}O_2$) | 102.1 | 5.0 | 26.0 |
| iC5/*i*-Valeric acid/*i*-valerate ($C_5H_{10}O_2$) | 102.1 | 5.0 | 26.0 |
| PentOH/Pentanol ($C_5H_{12}O$) | 88.10 | 5.0 | 30.0 |
| C6/Caproic acid/caproate ($C_6H_{12}O_2$) | 116.1 | 6.0 | 32.0 |
| iC6/*i*-Caproic acid/*i*-caproate ($C_6H_{12}O_2$) | 116.1 | 6.0 | 32.0 |
| HexOH/Hexanol ($C_6H_{14}O$) | 102.1 | 6.0 | 36.0 |
| C7/Heptanoic acid/heptanoate ($C_7H_{14}O_2$) | 130.1 | 7.0 | 38.0 |
| C8/Caprylic acid/caprylate ($C_8H_{16}O_2$) | 144.1 | 8.0 | 44.0 |
| $H_2$ | 2.016 | 0.0 | 2.0 |
| $CO_2$ | 44.01 | 1.0 | 0.0 |
| $O_2$ | 32.00 | 0.0 | -4.0 |
| $CH_4$ | 16.04 | 1.0 | 8.0 |
| CO | 28.01 | 1.0 | 2.0 |

**Table S4.** Carboxylate extraction efficiencies and selling prices assumed to estimate the value of broth.

| *Carboxylate* | *Extraction efficiency* (%) | *Selling price* (USD $kg^{-1}$) |
|---|---|---|
| C2 | 0.0% | 0.50 |
| C3 | 0.0% | 1.25 |
| C4 | 13.0% | 1.50 |
| C5 | 49.0% | 3.00 |
| C6 | 84.9% | 2.50 |
| C7 | 91.0% | 3.00 |
| C8 | 97.0% | 2.50 |
| iC4 | 13.0% | 1.50 |
| iC5 | 49.0% | 3.00 |
| iC6 | 84.9% | 2.50 |

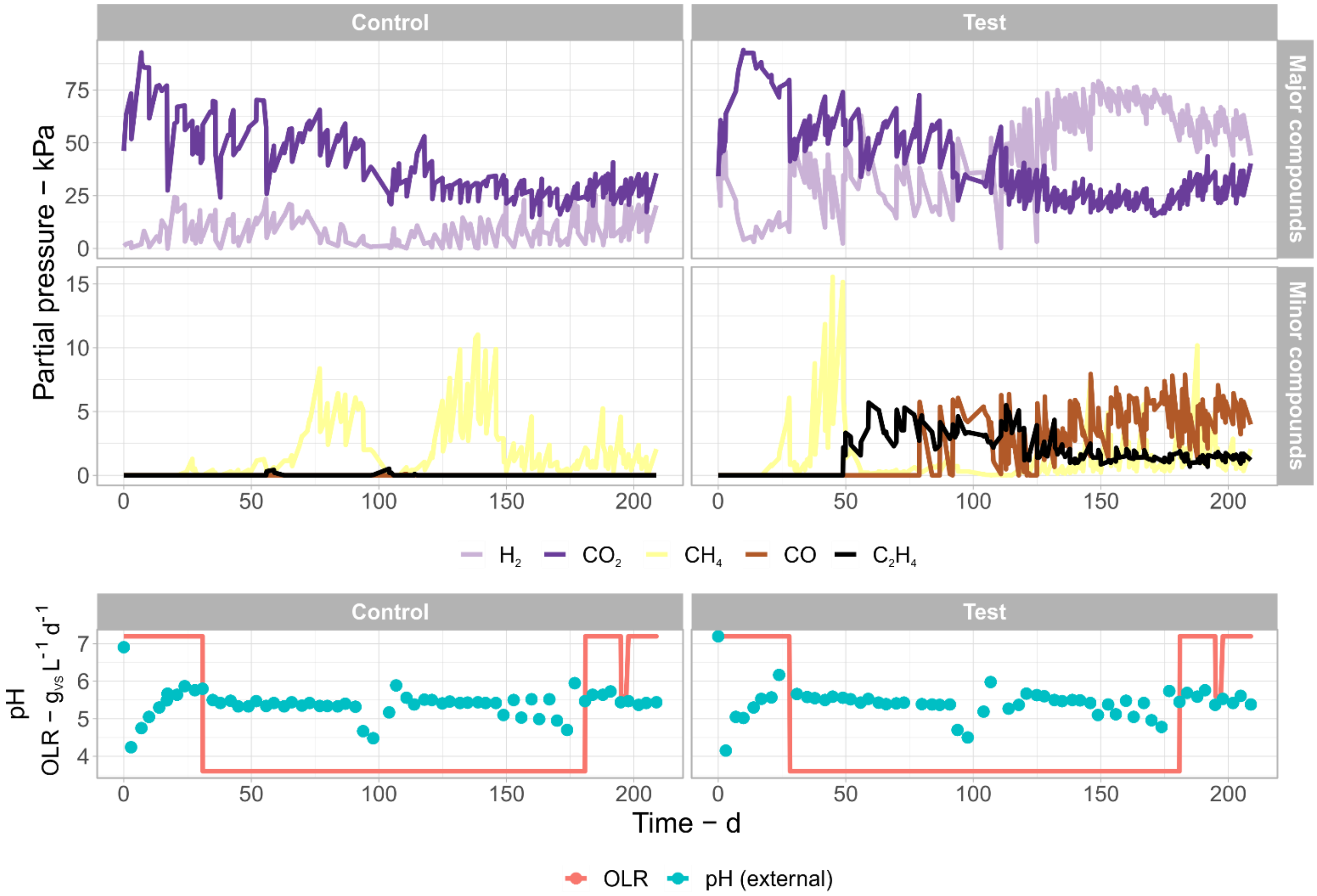

**Figure S1.** Partial pressures of gases, pH (measured externally), and organic loading rates (OLR) across the entire experiment.

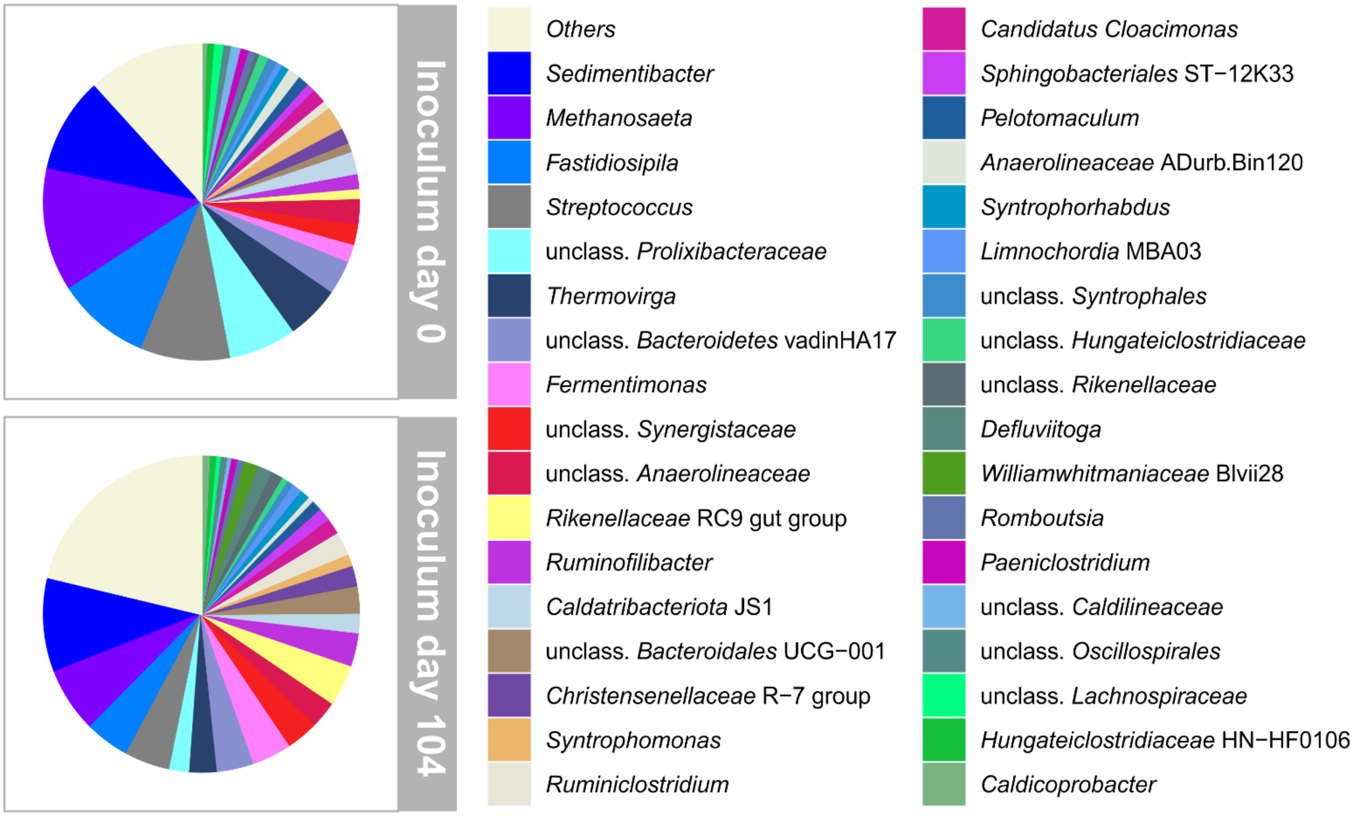

**Figure S2.** Microbial composition of inocula from days 0 (reactor startup) and 104 (reactor restart). Only the 35 most abundant genera in the inocula are shown.

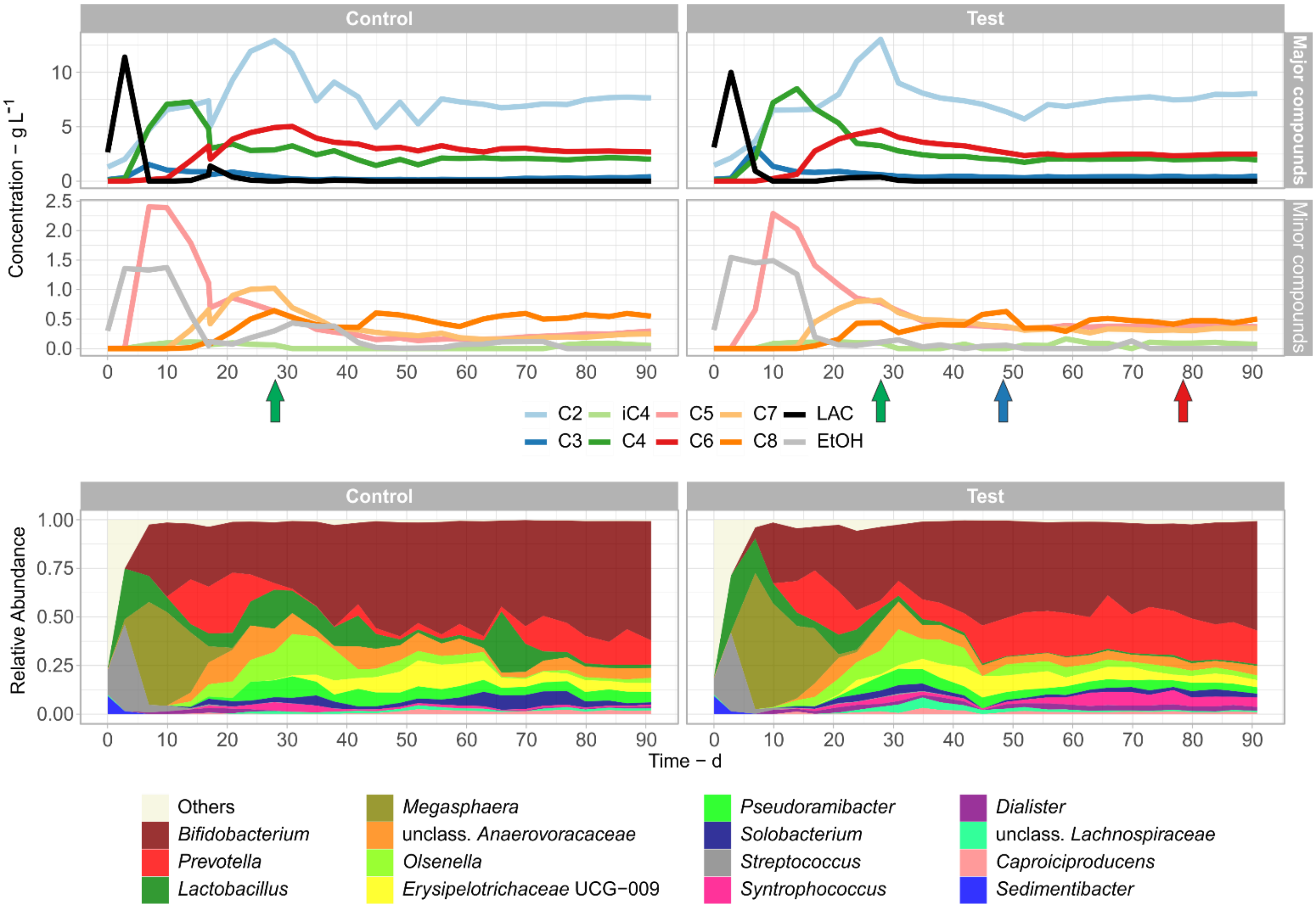

**Figure S3.** Chemical concentration and community composition during the first experimental period between days 0 and 91. The green arrows indicate a decrease of OLR. The blue and red arrows indicate the start of use of $C_2H_4$ and CO, respectively. Only the 15 most abundant genera are shown.

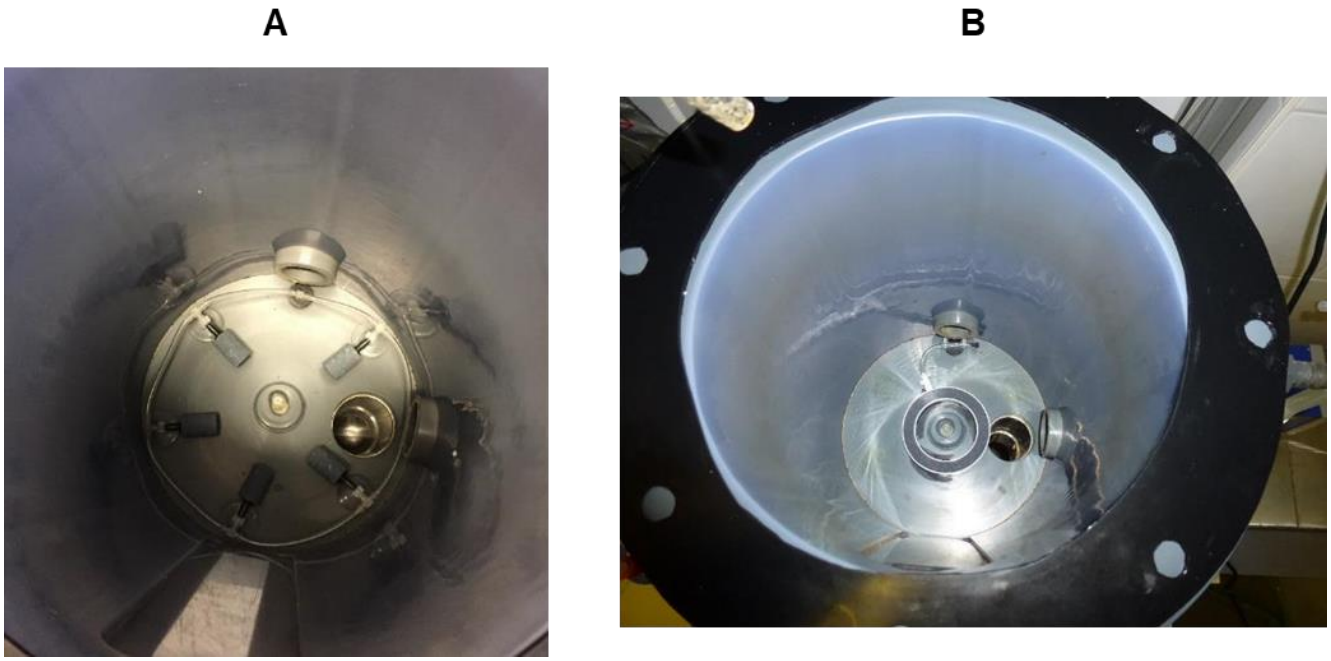

**Figure S4.** Two configurations of gas sparging. Firstly, multiple small spargers connected with PVC tubing (A) were used until day 98 when the spargers became dislodged. A ring-shaped sparger adhered to the bottom of the reactor (B) was used from day 104 until the end of reactor operation.

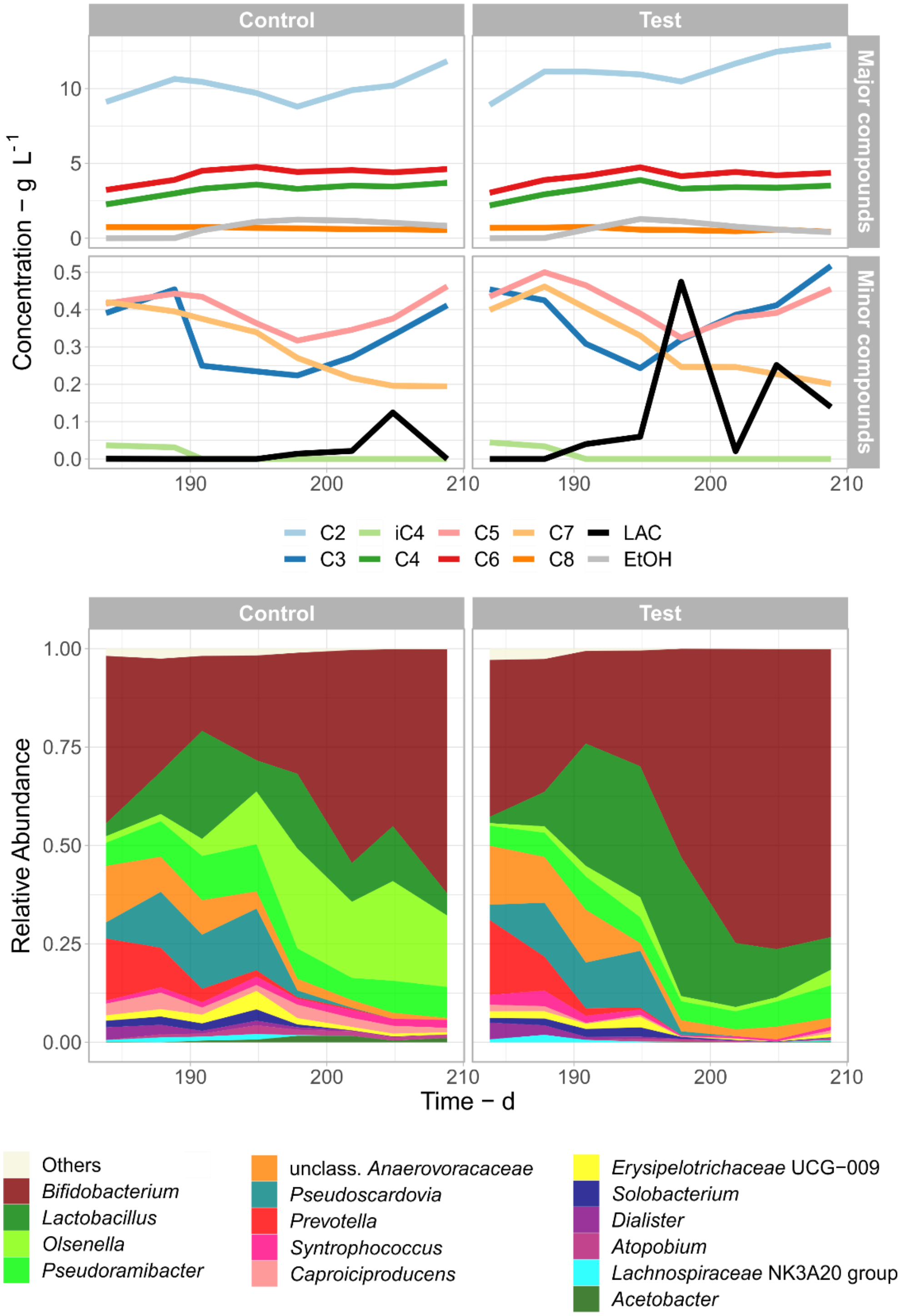

**Figure S5.** Concentration of chemicals and community composition between days 180 and 208. OLR was held at a high value (7.2 g VS $L^{-1}d^{-1}$). Only the 15 most abundant genera are shown.

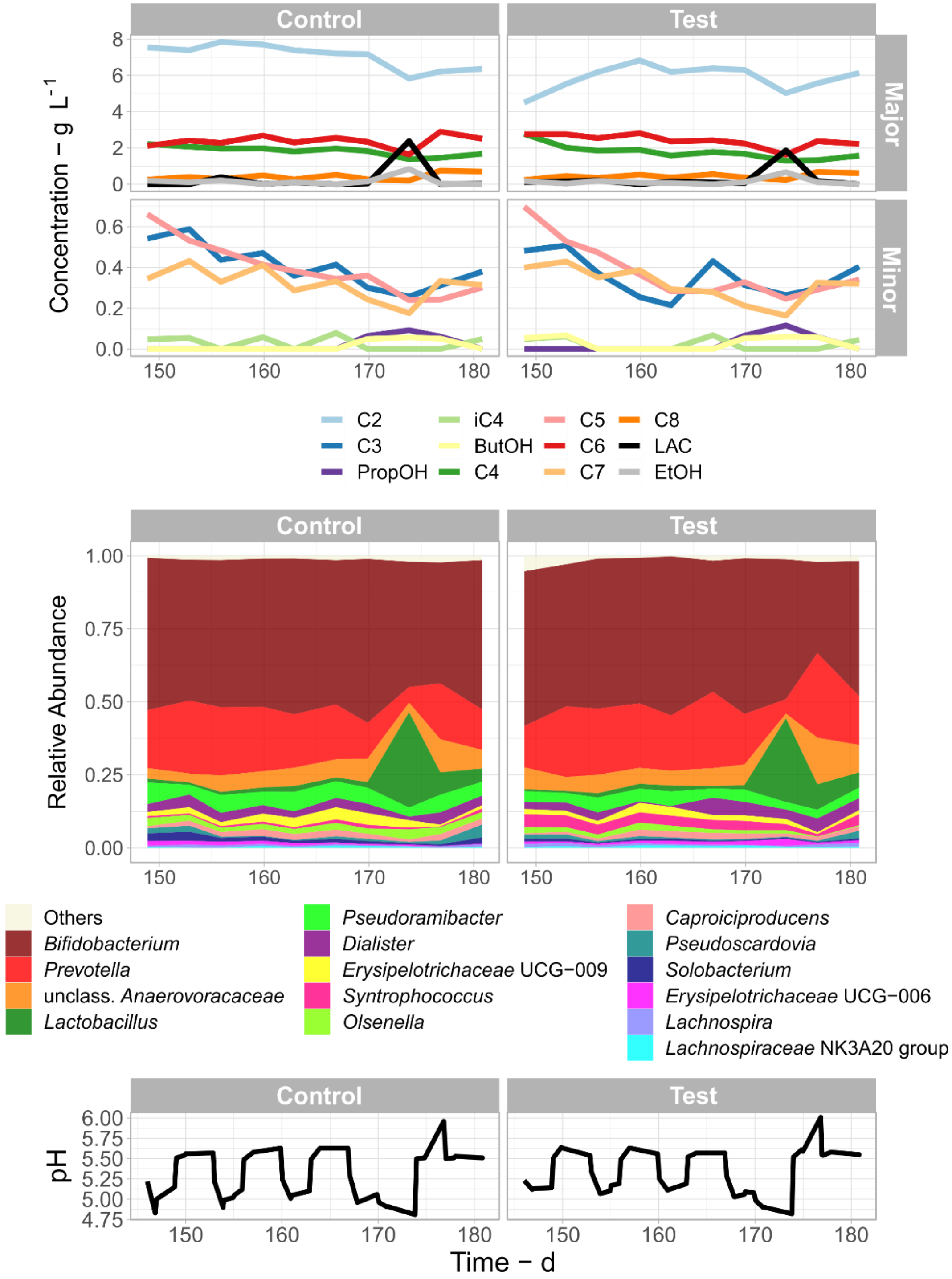

**Figure S6.** Concentration of chemicals, community composition, and pH between days 146 and 180. pH was varied in an attempt to induce solventogenesis. OLR was held at 3.6 g VS $L^{-1}d^{-1}$. Only the 15 most abundant genera are shown.

BIOREACTOR OPERATION AND GAS RECIRCULATION. pH was maintained using 10 M sodium hydroxide using a peristaltic pump controlled by a data acquisition and control unit (SENSOcontrol, Umwelt- und Ingenieurtechnik GmbH, Germany). The reactors were fed with corn silage, 3.5 mL of trace elements,[1] and 29.4 g $NH_4Cl$ twice a week. Corn silage was stored at 4°C in a plastic barrel, which was always flushed with $N_2$ for 10 minutes before storage. Two batches of corn silage were used throughout reactor operation, and their characteristics are reported in the Table S2.

Between days 98-104, fermentation in both reactors was stopped to replace the gas spargers, which had come out during reactor sampling. On day 98, the broth of each reactor was stored separately in opaque plastic barrels with $N_2$ in the headspace at 4℃ until the reactors were restarted. On day 104, reactors were restarted and inoculated using the anaerobically stored broth from day 98. CO and $C_2H_4$ were immediately added to the test reactor after this restart to match conditions before the reactor restart (Table S1).

The rest of the reactor system was isolated from the atmosphere while the gas bags were removed and refilled. Helium (He, 480 mL, >1% in the final mixture) was injected after each gas change and was used as an inert tracer to calculate material balances. $C_2H_4$ (480 mL, >1% in the final mixture) was injected into the test reactor to inhibit methanogenesis after day 49. The real gas composition available to the microbes differed from the gas bag filling ratios due to He and $C_2H_4$ addition, gas consumption, formation, and dilution with the reactor headspace.

CHEMICAL ANALYSIS. The ester-GC method described in Apelt [2] had the following modifications: a derivatization step by esterification using methanol was added and sulfuric acid was added instead of using phosphoric acid. The process for HPLC analysis was adapted from Apelt [3] is described as follows: remaining broth was filtered through a kitchen sieve to remove large particles and three sub-samples were centrifuged at 20,000 × g for 10 minutes. The supernatant of each sample was removed and passed through a 0.22 um nylon filter before being analyzed by high-performance liquid chromatography (Shimadzu). HPLC was performed using a refractive-index detector and a Hi-Plex H column (Agilent). The system was operated isocratically using a 0.005 M sulfuric acid solution as the mobile phase and a 20 μL sample injection volume. Column temperature was held at 55°C and flow rate at 0.7 $mL\text{-}min^{-1}$. The length of time for each sample was 180 minutes.

REACTOR STARTUP. During reactor startup (days 0-70), reactors were operated with $N_2$:$CO_2$ or $H_2$:$CO_2$. No CO was added to either reactor. Chemical concentrations were similar between the control and test reactor during this phase (Figure S3) and automatic pH control with setpoint at 5.5 only started operating effectively after day 10 (Figure S1). Following an initial spike in lactate, an increase in C3 and C4 was observed, with peak C4 concentrations of 7.2 g $L^{-1}$ and 8.5 g $L^{-1}$ in the control and test reactors, respectively. The rise and fall of C3 and C4 coincided with an increase and subsequent washout of *Megasphaera* (Figure S3) corresponding to its known ability to convert lactate into C3 and C4.[4] Similarly, spikes in lactate corresponded to peak abundances of *Lactobacillus* and *Streptococcus*. After day 10, despite the abundance of *Prevotella*, which is also a known lactate and C3 producer capable of breaking down cellulose and sugars,[5] concentrations of lactate remained near zero and those of C3 decreased quickly. Lactate is a valuable substrate for many acidogenic bacteria and is therefore expected to be present in very low concentrations during steady-state. The fall in C3 concentrations, on the other side, may imply that its production was mostly related to *Megasphaera* or that C3 was continued to be produced but it was converted into its elongated counterparts C5 (peak concentrations of about 2.3 g $L^{-1}$ in both reactors on day 10) and C7 (peak concentrations of about 0.9 g $L^{-1}$ in both reactors at around day

28). Expectedly, after OLR was reduced to 3.6 g VS $L^{-1}$ $d^{-1}$ on day 29, concentrations of all carboxylates decreased, in particular those of C6 and C2 (Figure S3).

EFFECTS OF MODULATING pH AND TEMPERATURE. To try to induce solventogenesis in the presence of abundant CO and $H_2$, pH and temperature were modulated every 2.8 or 6.8 days between 32°C/pH 5.5 or 28°C/pH 4.8. Production of PropOH and ButOH were used to assess whether syngas-triggered solventogenesis had occurred. EtOH was not used as an indicator since it is a common by-product of heterofermentative lactic-acid bacteria.[6] Although alcohol production by microbial communities fed with syngas was observed in other batch experiments,[7] in this experiment, measured PropOH and ButOH concentrations were minimal (<0.2 g $L^{-1}$).

As a result of changing the reactor conditions, in the test reactor, average $H_2$ consumption decreased from 9.35 to 1.71 mmol $L^{-1}$ $d^{-1}$ and average CO consumption decreased from 0.88 to 0.64 mmol $L^{-1}$ $d^{-1}$. $CH_4$ production in the test reactor, which had been minimal prior to the pH and temperature shift, increased from 0.16 to 0.66 mmol $L^{-1}$ $d^{-1}$. In contrast, $CH_4$ production decreased in the control from 0.79 to 0.38 mmol $L^{-1}d^{-1}$. A washout of *Megasphaera* during the pH cycling experiment (Figure S6) correlated with a decrease in C4 and increase in C2, further highlighting the role of *Megasphaera* in C4 production and the inability of other chain elongators present to fill this niche, despite *Prevotella* (023) showing similar correlations as *Megasphaera* (Figure 4). Possibly, *Megasphaera* is sensitive to pH and temperature, which could be used to fine-tune *Megasphaera* abundance and related C4 formation. CO did not appear to be able to overcome the pH and temperature effects at this partial pressure (5 kPa).

TEA 1 – ESTIMATING OPERATING COSTS AND CARBON EMISSIONS FROM GAS RECIRCULATION. The gas pressure increase expected from the compressor in an industrial setup together with the gas recirculation rate were identified as the two main factors influencing electricity consumption. Gas pressure increase depends on the plant detailed engineering and specific design aspects of the bioreactor such as sparging, tubing, and gas reservoir. In our analysis, we assumed a range of pressure increase values from 70 mbar (used in this study) to 2,000 mbar. The upper pressure limit is set so since high-pressure mixotrophic fermentation has never been tested. Moreover, high pressures incur in more expensive construction materials, which might easily set off economic advantages of the mixotrophic fermentation.

A constant gas composition of 0.75:0.185:0.05:0.015 ($H_2$:$CO_2$:CO:$C_2H_4$) with an average molar weight of 11.47 g $mol^{-1}$ at 20°C was assumed for calculating the electricity consumption of an isentropic compressor operating with 90% efficiency and pressure increases of 70, 500, 1000, and 2000 mbar. The average global electricity price in Q1 2025 of 0.15 USD $kWh^{-1}$ was assumed.[8] An effect of decreased stirring power consumption from gas sparging has not been accounted for as the gas flow rates assumed here are intrinsically low in comparison to aerobic fermentation setups, which typically aerate in the order of magnitude around of 1 vvm.[9]

The global average $CO_2$ electricity emission intensity in 2023 of 0.481 kg $CO_{2\,eq.}$ $kWh^{-1}$ [10] was adopted to calculate carbon emissions associated to electricity consumption.

TEA 2 – ESTIMATING THE COSTS OF GAS SUPPLY AND BROTH VALUE. A price of 0.264 USD $m^{-3}$ for acquiring carbon-neutral $H_2$ [11] was adopted for setting the $H_2$ + CO costs in Figure 5A. CO is not as commoditized as $H_2$, and, in an industrial setup, it will likely have to be generated from $H_2$ in-situ with a 1:1 reaction stoichiometry. For simplification purposes, the CO cost was assumed to be the same as that of $H_2$ on a volume basis (0.264 USD $m^{-3}$). A microbial $H_2$ + CO consumption rate of 252 mL $d^{-1}$ $L^{-1}$ (8.7% CO) with an HRT of 10 d was assumed to calculate the $H_2$ + CO acquisition rate.

For scenarios 1, 2, and 3 (Figure 6) costs for acquiring $H_2$ and CO were assumed as 0.132, 0.264, and 0.440 USD $m^{-3}$, respectively. Scenario 1 has a compressor increasing pressure by 500 mbar, in scenario 2 this value is 1000 mbar and in scenario 3, 2000 mbar. Adequate gas supply is achieved by only 0.5 vvh in scenario 1, whereas 1.0 vvh is necessary in scenarios 2 and 3. All three scenarios assume the same electricity cost (0.15 USD $kWh^{-1}$).

A compensation of 76.90 USD $t^{-1}$ $CO_{2\,eq.}$ [12] was assumed for calculating carbon credits in Figure 6.

To assess the value of the fermentation broth due to the more easily extractable carboxylates, an extraction efficiencies for different carboxylates based on the separation process developed by Braune, Yuan, Sträuber, McDowall, Nitzsche and Gröngroft [13] was assumed (Table S4). Assumed selling prices of carboxylates are also shown in Table S4.

Carboxylate concentrations were retrieved from the average carboxylate concentrations in the periods of “7 - low OLR” and “16 - high OLR” available in the spreadsheet made available as part of the Supporting Information.
